# TET2-mediated epigenetic modification promotes stress senescence of pancreatic β cells in type 2 diabetes mellitus

**DOI:** 10.1101/2024.11.01.621468

**Authors:** Weijuan Cai, Qingqing Song, Xiaoqing Mo, Huaqian Li, Yuling Song, Liang Yin

**Affiliations:** Institute of Clinical Medicine, Central People’s Hospital of Zhanjiang, Zhanjiang, 524000, China; Department of Endocrinology and Metabolism, Central People’s Hospital of Zhanjiang, Zhanjiang, 524000, China

**Keywords:** type 2 diabetes mellitus, cell senescence, DNA methylation, PTEN, H4K16ac

## Abstract

Epigenetic modification plays a key role in β cell senescence. In the regulation of gene expression, there is a complex and close relationship between DNA methylation and histone modification. In order to explore its specific mechanism in T2DM β cell senescence, we used postbisulfite aptamer labeling of genome-wide bisulfite-SEQ, chromatin immunocoprecipitation-SEQ, RNA-SEQ, CRISPR/Cas9 TETs knockout, RNA interference, TET2 inhibitors, lentiviral overexpression, and gene knockout mouse models. Our study found that demethylase TET2 was localized in the islets of mice, and the expression level increased with age. TET2 knockout in pancreatic β cells can hypermethylate PTEN, up-regulate MOF and enrich H4K16ac, and reduce the level of aging markers. This study confirmed that TET2-mediated PTEN DNA methylation can promote a new mechanism of β cell senescence by regulating H4K16ac, providing a new molecular mechanism and therapeutic target for T2DM β cell senescence therapy.

## INTRODUCTION

Type 2 diabetes mellitus (T2DM) is a growing global public health problem whose prevalence increases with age. T2DM in old age is usually accompanied by progressive impairment of β cell function, and β cell mass decreases with aging. However, the contribution of beta cell senescence to the pathogenesis of type 2 diabetes remains uncertain. Therefore, it is of great clinical value and social significance to explore the mechanism of aging of T2DM islet beta cells and seek new targets for the development of effective therapeutic drugs.

Epigenetic modification plays a key role in the aging of pancreatic β cells (Cagan et al., 2022). Epigenetics refers to the genetic mechanism that controls gene expression without affecting gene sequence in the regulation of gene expression (Barres & Zierath, 2016). It consists of various chromatin modifications, including DNA methylation, histone acetylation, and RNA modifications, which regulate cell differentiation, cell-specific gene expression, parental imprinting, X chromosome inactivation, and genome stability. Therefore, it is of great significance to explore the regulatory mechanism of β cell senescence from the perspective of epigenetics.

DNA methylation is one of the most widely studied epigenetic mechanisms. Changes in DNA methylation patterns related to β cell function (Dayeh et al., 2014) and identity have been shown in islet of diabetic patients and have been shown to affect β cell senescence and functional maturation (Avrahami et al., 2015; Golson & Kaestner, 2017), but little is known about the role of DNA demethylation in β cell senescence. The Ten-eleven translocation (TET) is located in the nucleus and oxidizes 5-methylcytosine (5mC) to 5-hydroxymethylcytosine (5hmC) for active DNA demethylation. TET proteins are Fe (II) and alpha-ketoglutarate-dependent enzymes, including TET1, TET2, and TET3. Studies have shown that loss of TET2 in pancreatic lineages leads to improved glucose tolerance and β cell function (Yang et al., 2018). Silencing TET2 can inhibit high glucose (HG) -induced levels of 5hmC and MMP-9 (Kowluru et al., 2016). Studies using zebrafish studies as a model further confirmed that HG activates TET, thereby inducing demethylation of cytosine throughout the genome (Dhliwayo et al., 2014). HG promotes the mRNA expression of TET2, which can lead to the dynamic changes of 5mC and 5hmC in white blood cells of T2DM patients (Yuan et al., 2019). In autoimmune T1DM individuals or during disease development, TET2 expression levels in human and mouse beta cells are significantly elevated due to inflammatory stimulation (Rui et al., 2021). In addition, studies have confirmed that the level of 5hmC in diabetic patients increases with age (Pinzon-Cortes et al., 2017). These studies suggest that TET2-mediated DNA demethylation may affect the function of islet beta cells. However, whether TET2 promotes islet beta cell senescence is not yet clear.

Our study intends to use post bisulfite aptamers to mark whole genome bisulfite-SEQ, chromatin immunocoprecipitation-SEQ, RNA-SEQ, CRISPR/Cas9 TETs knockout, RNA interference, TET2 inhibitors, lentivirus overexpression, islet beta cell-specific TET2 transgene and gene knockout mouse models, etc. From the cellular and animal levels, we elucidate the molecular mechanism of SGLT2i’s regulation of TET2-mediated PTEN hydroxymethylation and H4K16ac modification through inhibiting ROS levels to delay stress senescence of islet beta cells, in order to provide theoretical basis for SGLT2i’s treatment of senescence of islet beta cells in T2DM.

## 2 MATERIALS AND METHODS

### 2.1 Animal

TETs+/− and TETs−/− C57BL/6J mice were generated by GemPharmatech using CRISPR/Cas9 technology. Animal SPF grade db/db and db/m mice supplied by GemPharmatech. By mating heterozygous db/m TET2+/− mice, db/db TET2−/− and db/db TET2+/+ mice are generated. The experimental mice were all male. They were fed with normal chow diet (NCD) or high-fat diet (HFD) and sterile water at 22°C and certain humidity (45% ∼ 55%). All animal work was carried out under the guidance of the Animal Research Ethics Committee of Zhanjiang Central People’s Hospital (KYSB-DW-2024039).

### 2.2 Data Preparation

We take “islet beta cells aging” and “the pancreas senescence” as keywords, from GEO (https://www.ncbi.nlm.nih.gov/geo/) mRNA expression data retrieval and download. Select and download GSE72753 and GSE68618 for differential gene expression analysis. All of the above raw data can be downloaded for free from the GEO database. These datasets met the following criteria: (1) the species was mouse; (2) Aged pancreatic β cells and control tissue samples; (3) The sample was repeated at least three times in the experiment.

### 2.3 Islet isolation and insulin secretion assay

The islets were isolated by perfusion with collagenase V. According to the experimental content, the samples to be tested and KRBH solution were prepared on the same day, and the culture solution containing NG (5.5 mM) or HG (25 mM) was added, islets of uniform size were added, the supernatant was incubated for 30 m in and discarded, and the prepared samples to be tested were added, and the supernatant was extracted after incubation for 30 m in, and insulin content was determined by using insulin reagent kit.

### 2.4 Knock down TET2/PTEN in INS-1β cells

shRNA sequences targeting TET2/PTEN were designed and cloned into the lentiviral eukaryotic expression vector pLKO.1-puro. The shRNA sequence was 5’-AccgatgtCCTTgTAGCAC-3’ (TET2). 5’-gttggcgagtgttttgtgaag-3’ (PTEN) produced lentiviral particles in HEK293T cells transfected with pLKO.1-TET2-shRNA or PlKO.1-pyrrole plasmids and viral packaging plasmids psPAX2 and pMD2. G. INS-1 cells were exposed to the virus for 48 hours and screened with purinomycin (2.5 μg/ml) for 7 days. The efficacy of TET2/PTEN knockdown was verified by Western blot analysis of INS-1 cells.

### 2.5 IPGTT and GSIS

For IPGTT, after overnight starvation, mice were injected intraperitoneally with 0.5 g/kg of D-glucose, and blood glucose concentration in tail vein blood was measured by glucometer at 0, 15, 30, 60, 90, and 120 min after the injection, respectively. For GSIS, mice were starved for 6 h, and then injected intraperitoneally with 0.75 U/ kg of recombinant human insulin, and blood glucose concentration in tail vein blood was measured at 0, 15, 30, 60, 90, and 120 min after the injection, respectively. The blood glucose concentration was measured at 0, 15, 30, 60, 90 and 120 min after the injection.

### 2.6 RNA sequencing

Pancreatic islets were isolated from 24-week-old male WT and KO mice. BasePair BioTechnologies performed RNA extraction and RNA expression profile analysis. Briefly, to find differentially expressed genes in the samples, DEseq software was used for differential analysis.

### 2.7 Morphometric assessment

A portion of pancreatic tissue was fixed in 4% paraformaldehyde for IF, IHC and HE staining after embedding and sectioning. Pancreatic islet cells were prepared and stained using a β-Gal staining kit (Beyotime Biotechnology, C0602).

### 2.8 Traditional bisulfite sequencing and TET-assisted bisulfite sequencing (TAB-seq)

Genomic DNA was extracted from HG-cultured β-cell lines and processed using the EZ-DNA Methylation Direct Kit (Zymo Research). The treated DNA was amplified by PCR and purified with a gel extraction kit (Qiagen) and finally sequenced using standard Sanger sequencing. TAB-seq was performed by glycosylating genomic DNA extracted from β-cells with T4 phage β-glucosyltransferase (NEB), followed by oxidation with TET2, and finally by treatment with sulfite.

### 2.9 PBAT library preparation

The PBAT library was constructed using approximately 5ng genomic DNA to ensure that the library was performed on the Illumina HiSeq 4000 sequencer. Regions with at least 20% difference in absolute methylation levels between the TET-KO and WT samples were defined as hypermethylated and hypomethylated regions. A promoter is defined as the −1.5 to +1.5 kb region of the TSS (transcription start site).

### 2.10 WGS

Genomic DNA was isolated from rapid-frozen cells using the DNeasy kit. The genomic DNA was fragmented at 500bp peak by the ultrasound generator Covalis, and the TruSeq DNA library preparation kit added the DNA adapter to the double-stranded DNA according to Illumina manufacturer’s instructions. BGI performs deep whole genome sequencing on the Illumina HiSeq×10 platform.

### 2.11 Chromatin immunoprecipitation (ChIP) assay

ChIP assay was performed according to the protocol of EZ ChIP chromatin immunoprecipitation kit. The cross-linked chromatin DNA was ultrasound treated, followed by immunoprecipitation with antibodies. Normal IgG was used as a negative control. After purification, qRT-PCR was used to quantify the enrichment of related genes.

### 2.12 ChIP-Seq

ChIP DNA was prepared for sequencing according to TruSeq DNA Sample Preparation guidelines. In short, purified DNA of 10-50 ng of chromatin immunoprecipitation is obtained from different numbers of cells depending on the antibody used for ChIP, splice ligated and PCR amplified according to the manufacturer’s protocol. The sequencing library is multiplexed and run on Illumina sequencers. Finally, reads were quality filtered according to Illumina pipeline. All ChIP-seq experiments were independently repeated twice.

### 2.13 Library preparation

The library was prepared using Illumina’s NEBNext Ultra II DNA Library preparation kit. The ChIP libraries were sequenced with paired end readings of 2×50 bp on the Illumina NovaSeq6000 sequencer. Processing of ChIP-seq data sets. The data was normalized as a log2-fold change of H3 immunoprecipitation. Generate a block diagram using DeepTools2 multiBigwigSummary scores. The Bioconductor package ChIP-Seeker110 is used to retrieve the nearest genes around the peak, annotate the genomic region of the peak, and retrieve peak feature annotations.

### 2.14 Statistical analysis

Data were analyzed and plotted by GraphPad Prism 9.0 and R 4.2.3 software. All samples were randomly assigned to the treatment group. All samples represent biological replicas. All statistical measurements are expressed as mean ± standard deviation. The unpaired *t*-test (two-tailed) was used to assess the statistical significance of an observed parameter between two experimental groups. For comparisons of more than two groups, the data were analyzed using the univariate analysis of variance (ANOVA) of the Tukey test. *P* <0.05 was considered statistically significant.

## 3 RESULTS

### 3.1 Age-related DEGs from pancreatic β cells

In addition, age-related DEGs analysis was performed in pancreatic β cells using a high-throughput gene expression GEO database. The blood included a mouse expression dataset (GSE772753 and GSE68618). The up-regulated and down-regulated genes of volcanoes were shown in eight datasets using ggplot2 (Figure 1a). In addition, the heat maps of the top 20 differentially expressed genes were shown in Figure 3a (*P* < 0.05), as well as the 10 differentially expressed genes with the greatest up and down-regulation in the 2 expression data sets, as shown in Figure 1c, d. Importantly, we found that TET2 expression was significantly elevated in older mouse pancreatic β cells

**FIGURE 1.**
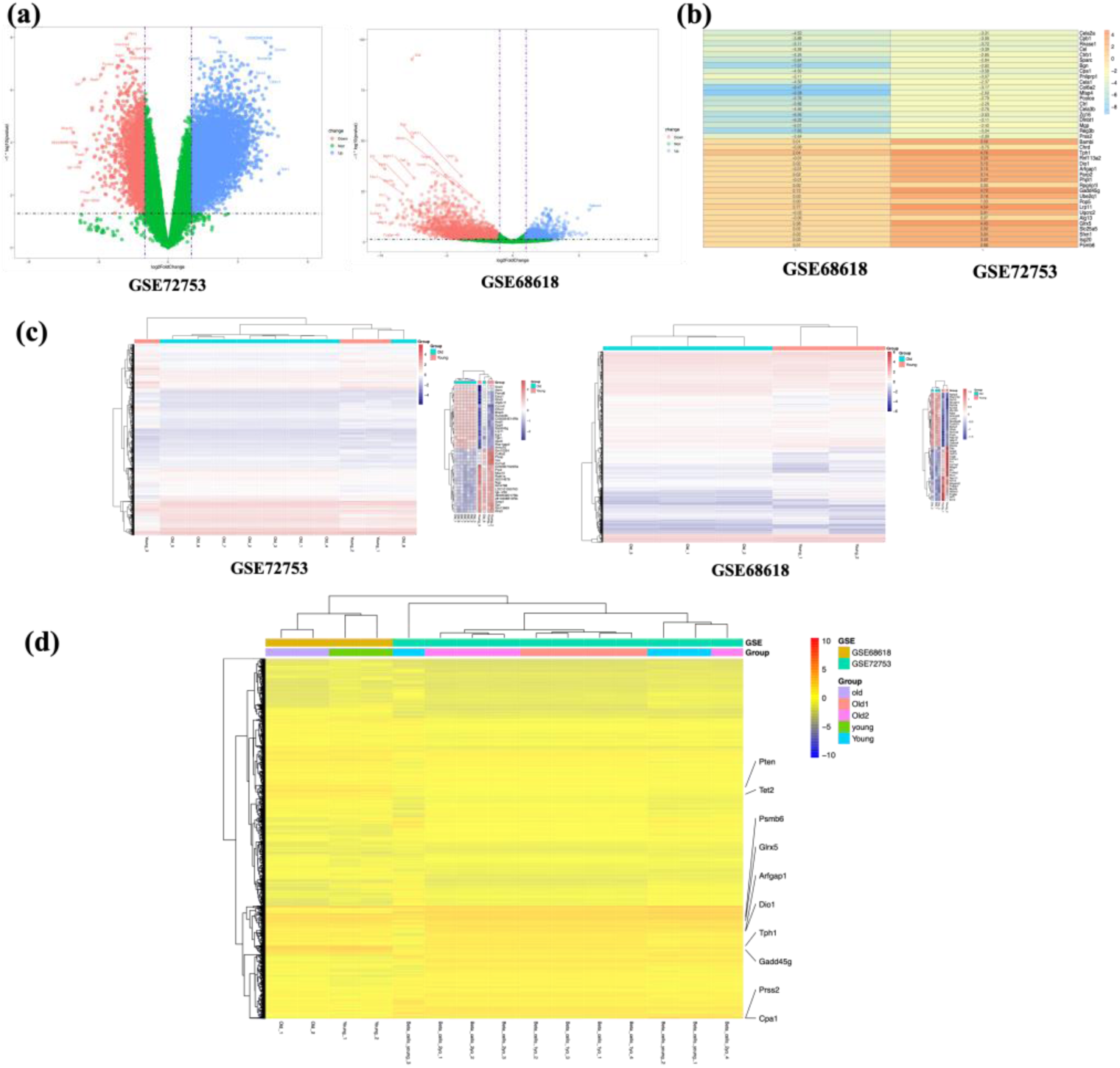
log2FC heat maps of age-related DEGs. (a) Volcano maps of DEGs in each dataset. (b) 2 Mus musculus expression microarrays in pancreatic β cells. (c) A representative image showing the heatmap, that on the right shows the top 10 DEGs between the older and younger groups. (d) Heatmaps of the differential gene expression for datasets. Red represents a high level of expression, blue represents a low level of expression, each column represents one sample and each row represents one gene. The specific locations of representative differential genes in the heat map have been marked on the right.

### 3.2 Functional enrichment analysis of differentially expressed genes in aging pancreatic β cells

Two gene expression profiles were identified by comprehensive transcription analysis and combined with GO analysis, 3218 functional items of 158 upregulated differentially expressed proteins were successfully annotated (Figure 2a). Biological processes (BP) include 2,283 functional items, the most important of which are biogenetic and metabolic processes, decomposition processes, and synthesis processes, Including proteasome-mediated ubiquitin-dependent protein catabolic process and ribonucleoprotein complex biogenesis, proteasome-mediated ubiquitin-dependent protein catabolic process, nucleotide metabolic process, and nucleotide biosynthetic process. We found 445 functional items in the Cellular Component (CC), These results indicated that differential proteins were mainly distributed in mitochondrial protein-containing complex, ribosome, transport vesicle and autophagosome. In addition, we found that molecular function (MF) accounted for 490 functional items, It mainly includes ubiquitin-like protein ligase binding, transcription coregulator activity and translation initiation factor activity, translation initiation factor binding and translation regulator activity, etc. When we analyzed the KEGG pathway, we found 52 significantly enriched up-regulated pathways, it mainly includes Parkinson disease, Protein processing in endoplasmic reticulum, Oxidative phosphorylation, Mitophagy and Ferroptosis (Figure 2b). Circos maps showed upregulated and down-regulated differentially expressed genes, and we also obtained interesting DNA demethylation and cell aging from functional concentrations, and found that TET2 and PTEN genes had such functional concentrations (Figure 2c, d).

**FIGURE 2.**
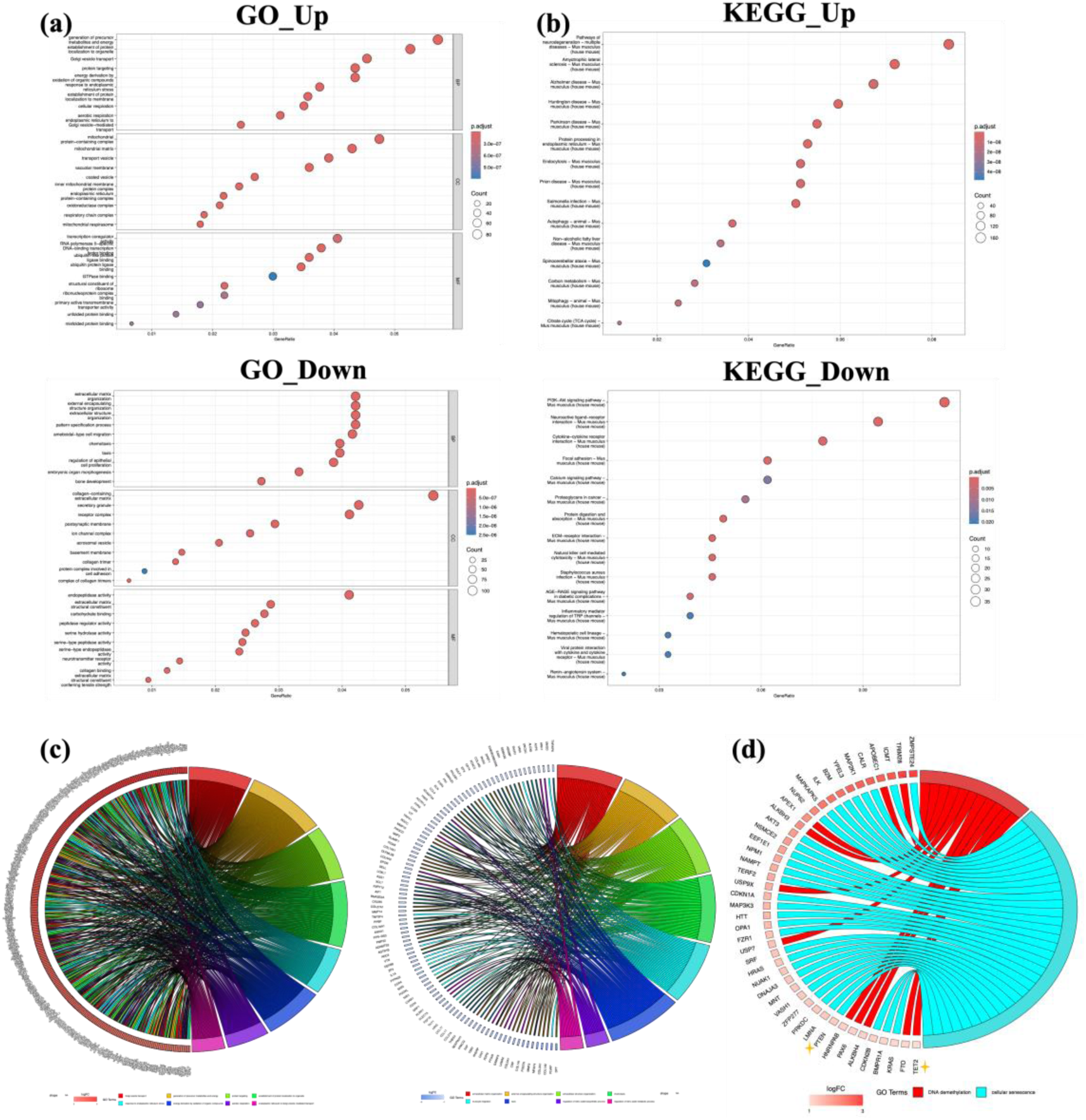
Functional enrichment analysis of aging islet β cells. (a, b) Enrichment analysis of differentially expressed genes GO and KEGG. The figure shows the GO and KEGG analysis results of mRNA up-regulation and down-regulation, respectively. (c, d) Circos maps show up-regulated and down-regulated differentially expressed genes (asterisks) that are key to GO enrichment in DNA demethylation and cell senescence.

### 3.3 TET2 is upregulated under conditions that promote diabetes

To determine the expression of TETs in beta cells, the rat beta cell lines INS-1E and MIN6 were exposed to an in vitro diabetic environment. TET2 protein levels are upregulated in both INS-1E and MIN6 cells (Figure 3a, b). Continuous feeding with HFD for 16 weeks detected a continuous increase in TET2 levels in the islets of the mice (Figure 3c). No significant difference in TET1 and TET3 protein levels was found between all groups, but TET2 protein levels were significantly higher in HFD than in the control group, and TET2 protein levels were also increased in the islets of db/db mice (Figure 3d). Dual staining of TET2 and insulin in paraffin-embedded sections of HFD confirmed β-cell-specific TET2 upregulation compared to NCD-fed mice (Figure 3e). TET2 expression was significantly increased in db/db beta cells compared to db/+ (Figure 1f). In addition, TET2 accumulates mainly in the nucleus (Figure 3g). The nuclear localization of TET2 is consistent with its DNA demethylation function. These data suggest that TET2 is significantly elevated in vitro and in vivo under pro-diabetic conditions.

**FIGURE 3.**
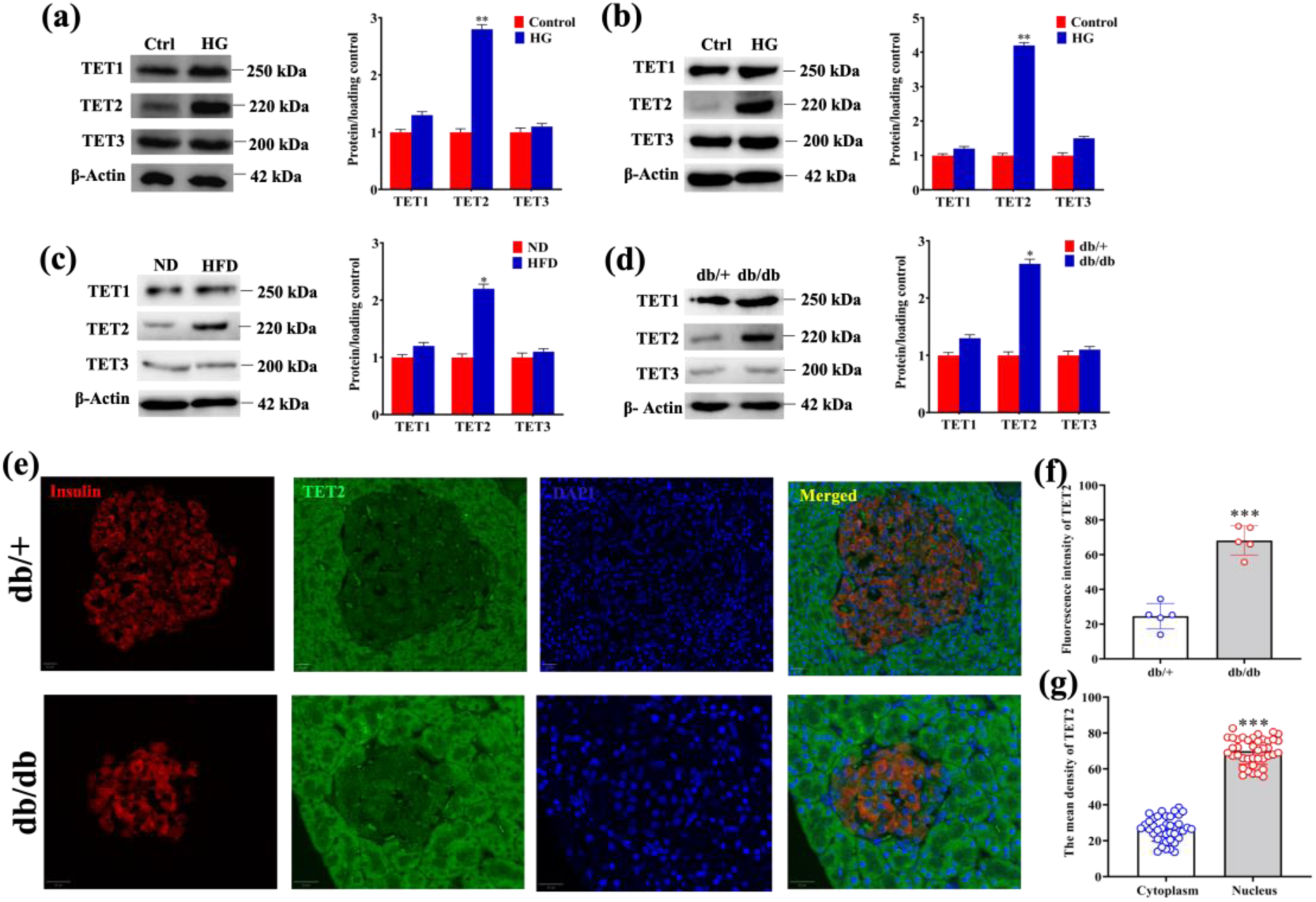
TET2 is upregulated in the occurrence of diabetes. (a, b) Western blot and analysis of INS-1E cells or MIN6 treated with high glucose (16.7 mM) for 2 days. (c) Western Blot and analysis of isolated islets in mice fed with NCD or HFD for 16 weeks. (d) Western blot analysis of isolated islets of diabetic db/db mice and their hybrid db/+ litters at 10 weeks of age. (e, f) IF sections of the db/+ and db/db mouse pancreas showed TET2 in red and insulin in green. (g) The average density of TET2 in the nucleus was higher than that in the cytoplasm.

### 3.4 TET2 loss improves age-related β cell function

To investigate the role of TET2 in glucose homeostasis, TET2 WT/KO mice were constructed and glucose tolerance tests were performed on WT and TET2 KO mice at 8, 16, 32 and 52 weeks, respectively (FIG. 4a). The results showed that the loss of TET2 improved the mice’s glucose tolerance from week 16 onwards, and the gap between WT and KO mice increased with age. At the same time, TET2 deficiency also promoted insulin secretion in mice (Figure 4b), and compared with WT mice, the islets of TET2 KO mice had higher levels of glucose-stimulated insulin secretion (Figure 4c). This enhancement may be responsible for the improved glucose tolerance in TET2 KO mice. In addition, we observed that TET2 expression was localized in the islets and increased with age (Figure 4d, e), suggesting that age-related TET2 loss may improve glucose tolerance to some extent. We treated WT and TET2 KO mice with HFD. It was found that HFD-treated TET2 KO mice also showed improved glucose tolerance with age (Figure 4f). TET2 is involved in glucose metabolism in an age-related manner. To confirm the specificity of TET2 in regulating insulin release and glucose metabolism, researchers conducted a series of experiments in TET1 and TET3 deficient mice, and found that WT, TET1 and TET3 KO mice had no significant differences in glucose tolerance, insulin release, etc. (Figure 4g, h), suggesting that with the increase of age, TET2 can specifically affect the function of islets and affect glucose metabolism in mice.

**FIGURE 4.**
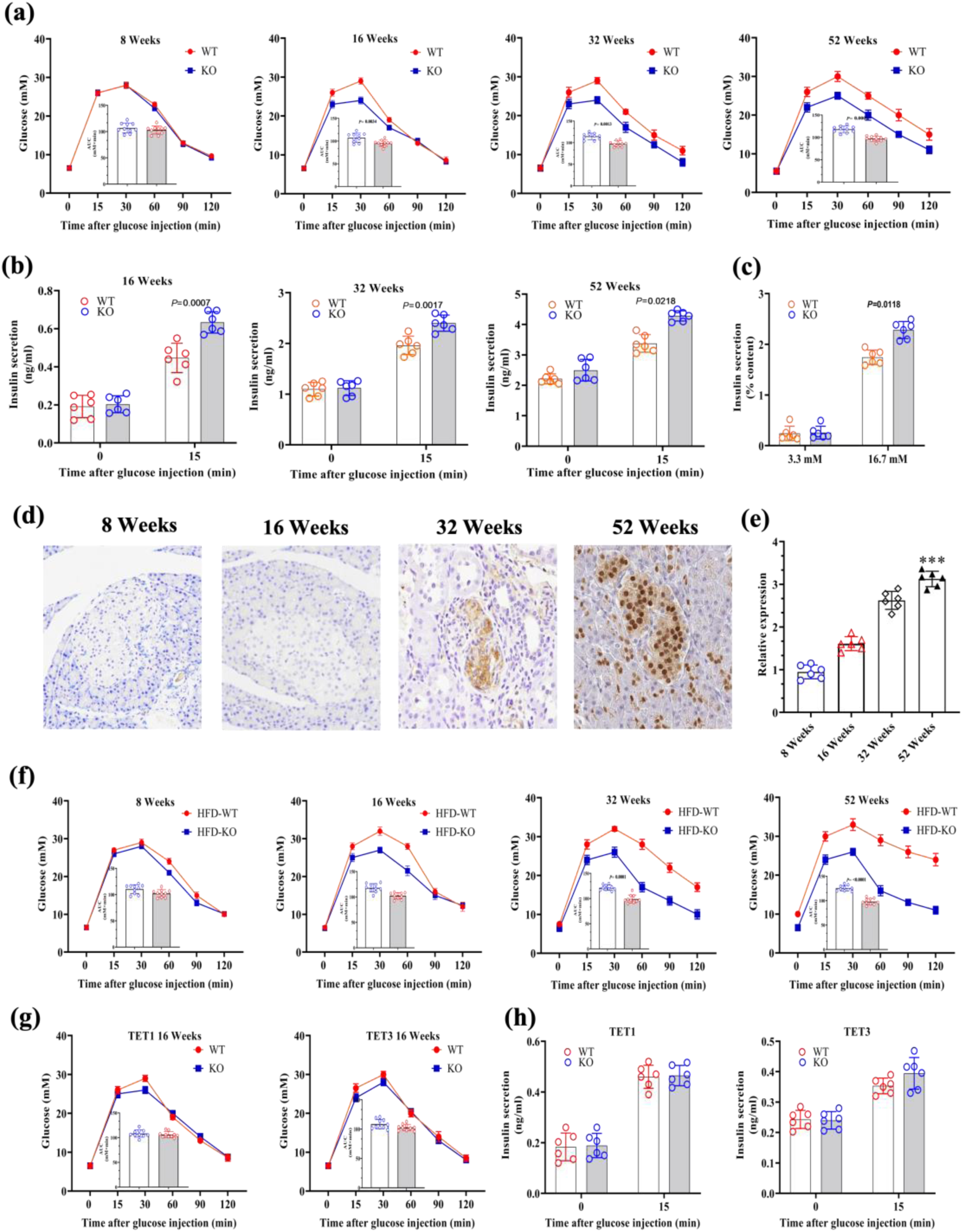
In NCD and HFD mice, TET2 KO mice showed improved glucose tolerance due to age-related increased insulin secretion. (a) IPGTT was performed in TET2 WT and KO mice fed NCD at weeks 8, 16, 32, and 52 to measure blood glucose concentration and area under the curve (AUC) of IPGTT. (b) Plasma insulin levels in TET2 WT and KO mice fed NCD at 16, 32, and 52 weeks. (c) Release of insulin from mouse islets isolated from 16-week-old TET2 WT and KO mice under low or high glucose conditions. (d) Representative IHC images of TET2 expression in the pancreas of WT mice at a set time point. (e) Statistical quantification of IHC images. (f) The IPGTT and AUC of 6-week-old male TET2 KO and WT mice were analyzed after HFD treatment. (g) NCD treated IPGTT and AUC in WT and TET1/TET3 KO mice at 16 weeks. (h) Insulin secretion in WT and TET1/TET3 KO mice was assessed at a specified time. P-values are displayed accurately.

### 3.5 TET2 loss can improve the aging and characteristics of beta cells

To explore the underlying mechanisms of improved glucose metabolism and increased insulin secretion in TET2-deficient mice, RNA sequencing was performed on islets obtained from 24-week NCD TET2-WT and KO mice and the expression of key β cell identity genes (including Mafa, Glut2, Nkx6-1, and Pdx1) was found to be enhanced in TEt2-deficient mice. However, the expression of age-related characteristic genes, including IGF1R, p16, and β-Gal, was reduced (Figure 5a), which was confirmed by QRT-PCR (Figure 5b). Immunofluorescence analysis was further used to detect increased expression levels of key β cell markers in the islets of TET2 KO mice under HFD conditions (Figure 5c). Previous studies have shown significant downregulation of key beta cell identity genes in senescent cells. The expression of aging markers in the islet beta cells was subsequently examined, and it was found that p16, IGF1R and β-Gal staining were significantly reduced in the islets of TET2-KO mice under HFD conditions compared with the WT control group (Figure 5d), indicating that TET2 loss can inhibit β cell aging. These findings suggest that TET2 loss inhibits aging-related beta cell senescence, contributing to better retention of beta cell properties.

**FIGURE 5.**
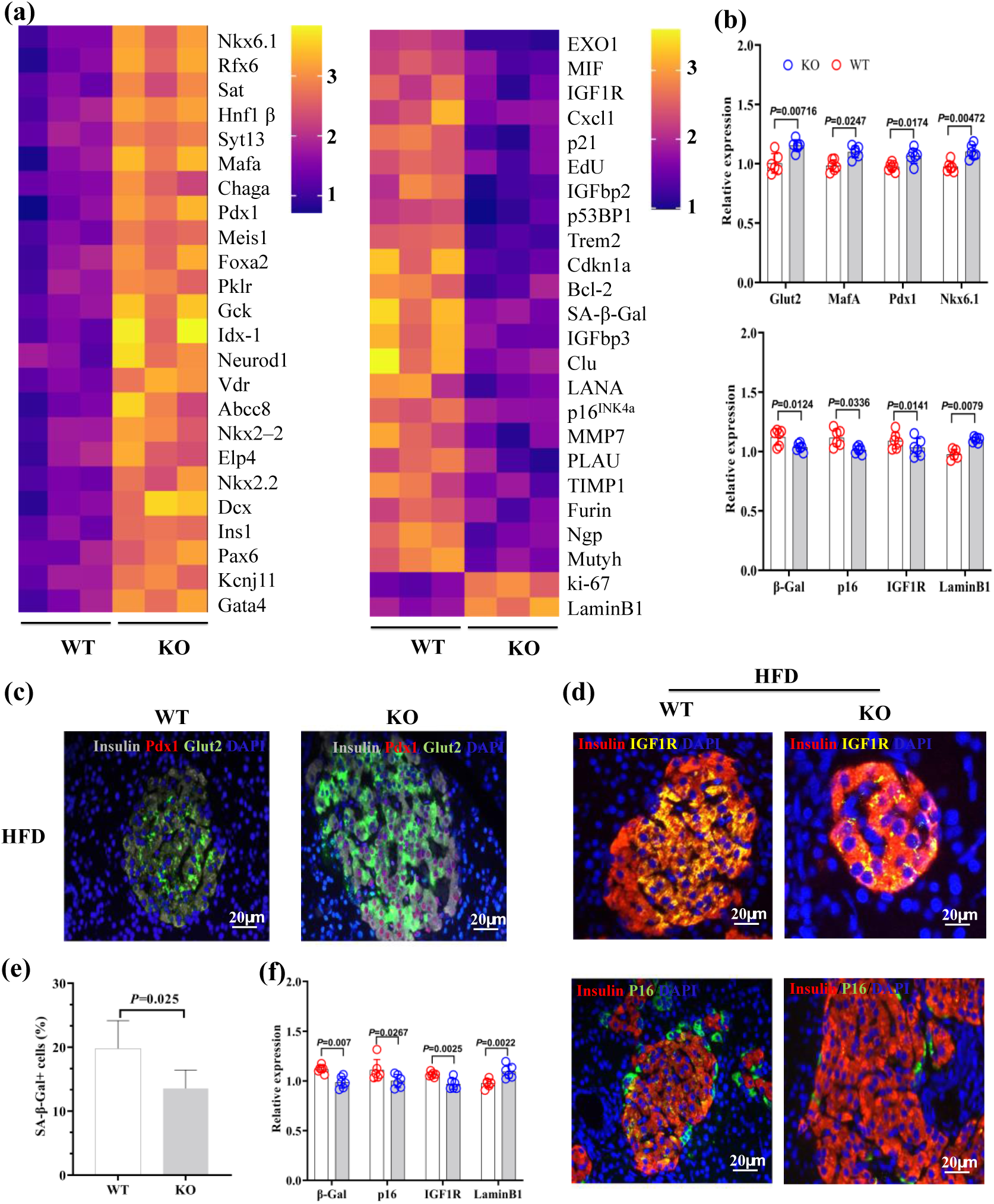
TET2 deletion enhances β-cell signature gene expression by antagonizing β-cell senescence in NCD or HFD mice. (a) The pancreatic islets of 24-week-old TET2 KO mice were treated with RNA-Seq, which showed up-regulation of β-cell characteristic genes and down-regulation of age-related characteristic genes (n= 3 per group). (b) QRT-PCR to verify the sequencing results. (c) Representative immunofluorescence of Pdx1 (red) or Glut2 (green) and insulin (gray) in islets of WT and KO mice treated with HFD (52 weeks). (d) Representative immunofluorescence of p16(green), IGF1R(yellow), and insulin (red). (e) Percentage of β-Gal staining positive in pancreatic islets of WT and KO mice treated with HFD. (f) Quantification of p16, β-Gal, IGF1R or LaminB1 immunofluorescence.

### 3.6 Overexpression of TET2 accelerated the aging of beta cells

To further analyze the specific effect of TET2 on β cell function, INS-1E cells were infected with LV-TET2 OE. Western blot confirmed overexpression of TET2 in INS-1E (Figure 6a). The results showed that TET2 OE decreased the expression level of β-cell identity genes and increased the expression level of aging markers (Figure 6b-d). The results suggest that TET2 plays a crucial role in regulating beta cell senescence.

**FIGURE 6.**
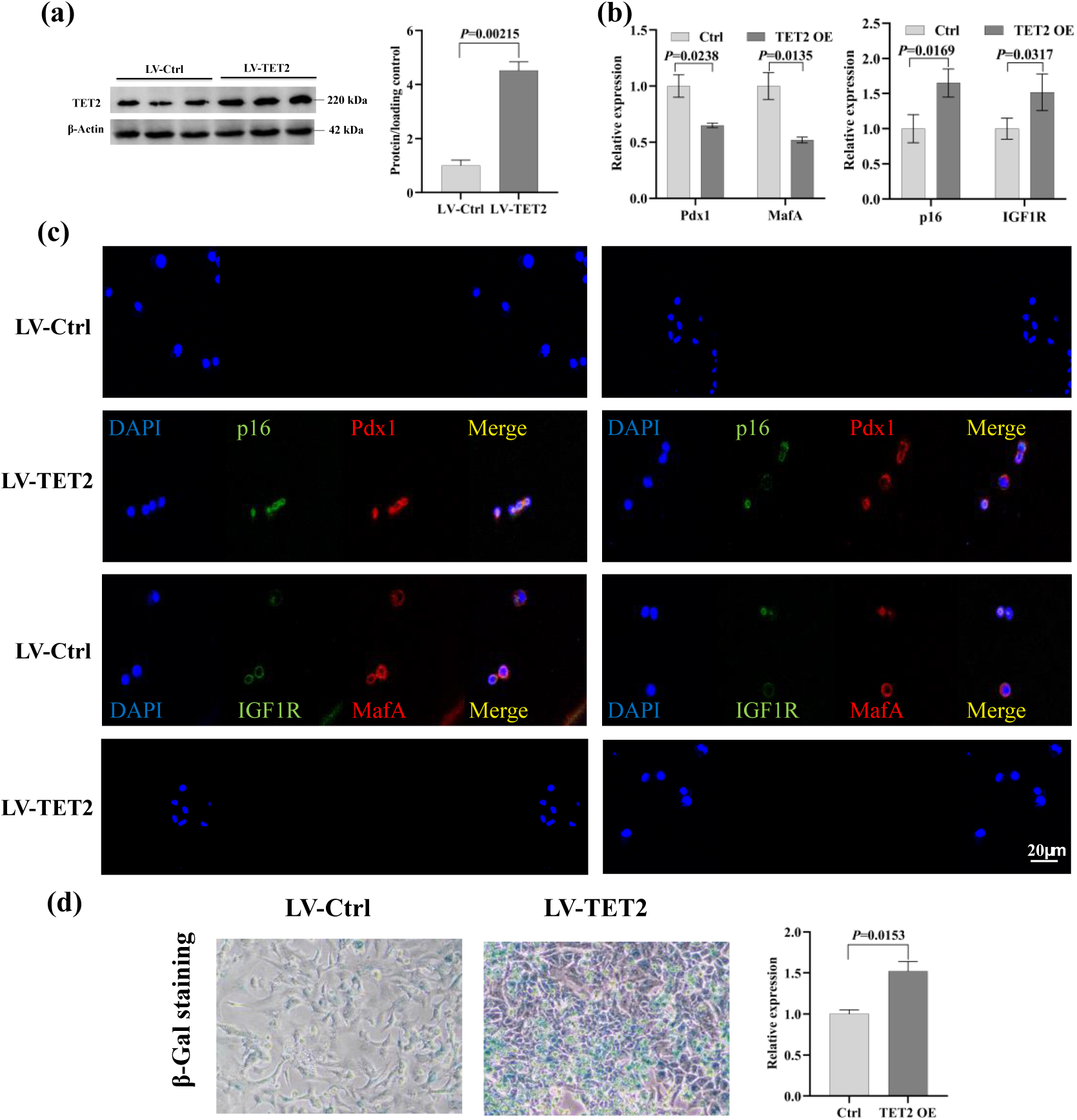
overexpression of TET2 accelerates the aging of β cells. (a-c) For MafA, p16 (red) and Pdx1, IGF1R (green). IF statistical quantification of MafA, Pdx1, p16 and IGF1R. (d) β-Gal staining and quantification.

### 3.7 Genome-wide bisulfite sequencing analyzed the effect of TET deletion on PTEN promoter methylation

To assess the overall effect of TET loss on DNA methylation, PBAT-WGBS was used in INS-1E. CpG methylation levels in TET-KO INS-1E cells (65.2%) were found to be comparable to those in WT INS-1E cells (68.8%) (Figure 7a). After dividing the PBAT signal into 100bp blocks, the TET-KO apparent cells showed a higher number of hypermethylated cells (69,513) than hypomethylated cells (50,327) (Figure 7b). Further analysis of various genomic features revealed that the hypermethylation sites in TET-KO INS-1E were mainly concentrated in promoters, enhancers, and DNase i hypersensitive sites (Figure 7c, d). In addition, using stricter cutoff values, 2,023 hypermethylated DMRs and only 39 hypomethylated DMRs were identified in TET2-KO INS-1E cells compared to WT, and the dominance of hypermethylated DMRs was as expected (Figure 7e). Gene ontology analysis of genes with hypermethylated enhancer and promoter regions in INS-1E revealed the enrichment of genes associated with developmental processes, including cell fate commitment, cell cycle, beta cell differentiation, and transcriptional regulation (Figure 7f). The data suggest that in TET-deficient islets, hypermethylated CpGs tend to cluster at specific genomic sites (promoters and enhancers) with gene regulatory potential. Gene-specific examination of the methylation group data showed that PTEN expression was significantly reduced in TET2-KO INS-1E and DNA methylation of its promoter was significantly increased compared to the control group (Figure 7g, h). Bisulfite Sanger sequencing confirmed ultra-low methylation of the PTEN promoter region in wild-type INS-1E cells (only 3% 5mC) and significantly increased methylation in TET2-KO INS-1E cells (27% 5mC) (Figure 7i). The PTEN promoter region was analyzed by TAB sequencing. Production of 5hmC in these regions can be saved by ectopic expression of TET2 in KO INS-1E (Figure 7j). Together, these data demonstrate that the inactivation of TET leads to elevated methylation of regulatory elements of the PTEN gene, which can lead to reduced expression and associated phenotypes.

**FIGURE 7.**
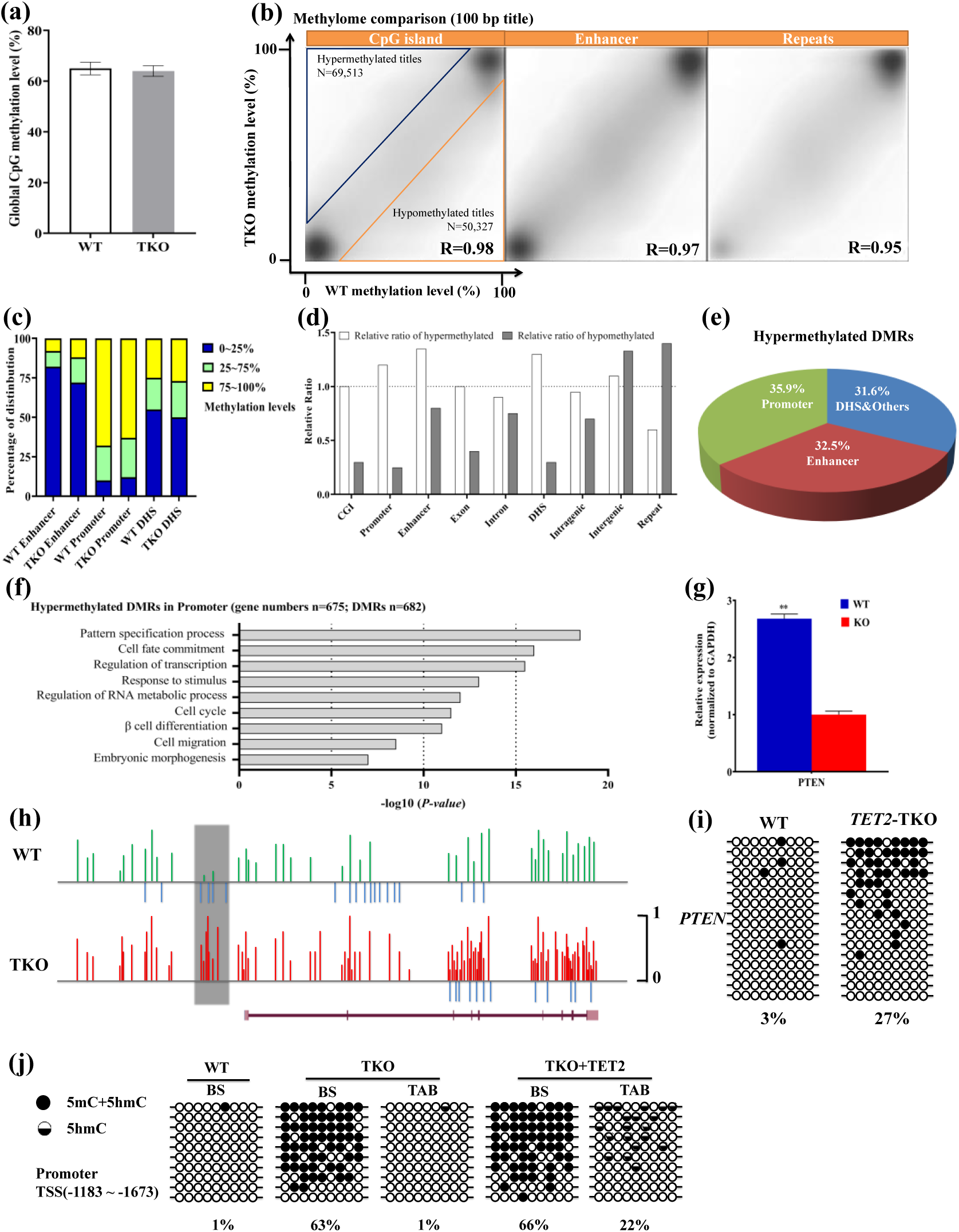
Methylation levels of PTEN promoter region in TET2 deficient INS-1E were analyzed by PBAT-WGBS. (a) Bar chart showing genome-wide CpG methylation levels for INS-1E wild type and KO. (b) Scatter plots showing a global methylation comparison between wild type and TET KO INS-1E using 100bp plots. Regions with at least 20% difference in absolute methylation levels between TET-KO and wild-type samples were defined as differentially methylated regions: hypermethylated regions and hypomethylated regions. The triangle represents a methylation difference of at least 20% between the wild-type and TET-KO samples. (c) Distribution of individual CpG methylation levels among the genomic elements shown. DHS: DNase I hypersensitive site. (d) Relative enrichment of hypermethylated and hypomethylated regions with different genomic features in INS-1E KO. CGI: CpG Island. (e) Pie chart showing the distribution of hypermethylated DMR in INS-1E KO apparent cells. (f) Gene ontology analysis of all genes with high methylDMR in the KO INS-1E promoter region. (g) Relative quantitative expression of PTEN in TET-KO INS-1E. (h) Methylation characteristics of the 5’region of PTEN gene in INS-1E extracted from PBAT whole genome sequencing data. The gray shaded box indicates the analysis area. Vertical bar above horizontal line (wild type, green; KO, red) indicates the methylation level (0-1) of a single CpG binary (counting two complementary CPGS), and the blue bar below the horizontal line indicates the detected unmethylation of CpG to distinguish it from the undetected CpG sites. (i) Methylation analysis of PTEN promoter in INS-1E by bisulfite-Sanger sequencing. The hollow and black circles represent unmethylated and methylated CpG sites, respectively. (j) TET assisted sodium bisulfite (TAB) sequencing analysis of PTEN promoter region.

### 3.8 Overexpression of TET2, but not TET1 and TET3, down-regulated the overall H4K16ac level

In the regulation of gene expression, there is a complex and close relationship between DNA methylation and histone modification. A comprehensive understanding of this interrelationship is lacking. Since TETs is the main demethylase, whether TETs is involved in the regulation of the overall level of H4K16ac remains to be studied. For this purpose, we performed H4K16ac ChIP-seq in INS-1E, MIN6, and βTC6 cell lines. H4K16ac was found to be a rich histone modification widely present throughout the genome (Figure 8a). In the tested cell lines, H4K16ac was significantly enriched on the genome, but significantly absent on TSS (Figure 8a-c). In addition, the high abundance of H4K16ac in INS-1E and MIN6 (approximately 30% of all histone H4) was confirmed using mass spectrometry (Figure 8d). To analyze the changes of H4K16ac after OE TETs, TETs OE was subjected to H4K16ac ChIP-seq in INS-1E cells. TET2 OE causes widespread deletion of H4K16ac on a genome-wide scale, except for TSS, which mostly maintains low signaling (Figure 8e-h). Surprisingly, TET1 and TET3 OE did not cause changes in their target gene H4K16ac, nor did they cause changes at the overall level (Figure 8e-h). Similar results were observed by Western blot analysis of overall H4K16ac levels (Figure 8i). In summary, these data suggest that TET2 down-regulates overall H4K16ac levels outside the TSS region in beta cell lines.

**FIGURE 8.** Overall level of H4K16ac elimination by overexpression of TET2 rather than TET1 and TET3. (a) H4K16ac-ChIP-seq signal heat maps of all genes in INS-1E, MIN6, and βTC6 cell lines. TSS, transcription start site. (b) Density maps of standardized MIN6 H4K16ac ChIP-seq signals in INS-1E cells for different chromatin states defined by the Broad Institute ChromHMM project. The colors in the density map convey a shape that is normalized to the maximum signal distribution within a channel. The quantity H4K16ac is marked on the Y-axis. (c) Map showing the distribution of statistical summaries of H4K16ac ChIP-seq signals in INS-1E cells grouped according to expression quantile (Q) distribution. (d) Bar plots showing the abundance of a single H4 peptide (amino acid 4-17) with different acetylation combinations in INS-1E and MIN6 cells as measured by mass spectrometry. (e) H4K16ac ChIP-seq signal heat maps of all genes in the TETs OE sequence. (f) Average normalized H4K16ac ChIP-seq spectra of all genes in the TETs OE sequence. (g) qRT-PCR quantized H4K16ac ChIP signaling at selected sites after OE TET2, TET1, or TET3. GB, the genome. (h) Examples of H4K16ac ChIP-seq tracks in the OE TETs series. (i) Western blot analysis of H4K16ac and H3 levels after OE TET2, TET1 or TET3.

### 3.9 TET2 promotes β cell senescence through the PTEN/MOF/H4K16ac axis

To explore the mechanism of TET2 in promoting β cell senescence, RNA-seq results of TET2−/− islets were analyzed, and MOF was significantly upregulated in TET2−/− mice (Figure 9a). This was confirmed by qRT-PCR (Figure 9b), suggesting that upregulation of MOF may contribute to better beta cell function in TET2-deficient mice. Further detection of MOF expression in INS-1E cells showed that MOF was inhibited when TET2 was overexpressed (Figure 9c), which was consistent with H4K16ac ChIP-seq results. Subsequently, the expression of PTEN and beta cell identity genes in TET2-OE INS-1E cells was analyzed, and PTEN was found to increase with the decrease of beta cell specific genes (Figure 9c). In addition, shPTEN reversed TET2 OE-induced changes in MOF and Pdx1 (Figure 9d). To further determine whether the MOF pathway is involved in the increase of β cell senescence, MOF expression was induced in INS-1E cells and p16 was significantly inhibited after MOF OE (Figure 9e). OE MOF further salvaged its expression in TET2-OE cells and found that the expression level of aging markers was significantly reduced, while the expression level of beta cell identity genes was increased (Figure 9f). These results were confirmed in INS-1E cells with double overexpression of MOF and TET2 (Figure 9g). In addition, TET2 knockdown in INS-1E cells confirmed that when TET2 is knocked down, the MOF signaling pathway is enhanced, the expression of aging markers is reduced, and insulin secretion is increased (Figure 9g, h). Taken together, these results suggest that TET2 regulates H4K16ac modification via the PTEN/MOF/ axis, thereby promoting β cell senescence.

**FIGURE 9.** TET2 regulates the PTEN/MOF/H4K16ac signaling pathway to influence β cell function. (a) Transcriptome heat maps of islets in mice aged 24 weeks (n=3). (b) qRT-PCR analysis in WT and KO islets. (c) Effects of TET2 OE on PTEN, β cell senescence and β cell recognition genes. (d) Effect of shPTEN on TET2 OE induced changes. (e) Expression of H4K16ac and p16 in INS-1E cells overexpressing MOFs. (f) MOF overexpression treatment for 24 h confirmed the effect of TET2 on β cell function through MOF. (g) Expression of p21 and p16 in INS-1E cells with TET2 overexpression or MOF overexpression or double overexpression. (h) Effects of TET2 knockdown on PTEN/MOF signaling pathway and aging markers in INS-1E cells. Ctrl, contrast; KD, knock it down. Effect of TET2 downregulation on insulin secretion under low and high glucose conditions.

## 4 DISCUSSION

Aiming at the breakthrough of the interaction between DNA methylation and histone acetylation, we carried out an in-depth experimental study. Firstly, in vitro and in vivo models determined TETs expression in beta cells, and results showed that TET2 was significantly upregulated in both pro-diabetic conditions. Then, to study the role of TET2 in glucose homeostasis, TET2 WT and TET2 KO mice were constructed, and the results showed that TET2 loss improved glucose tolerance in mice, and the gap between WT and KO mice increased with age. At the same time, TET2 deficiency also promoted the secretion of insulin in mice. In addition, it was observed that TET2 expression was localized in the islets and increased with age, suggesting that age-related TET2 loss may improve glucose tolerance to some extent. Subsequently, TET2 WT and KO mice were fed a High-fat diet (HFD) and found that HFD-treated TET2 KO mice also showed improved glucose tolerance with age, suggesting that TET2 is involved in glucose metabolism in an age-related manner. To further confirm the specificity of TET2 in regulating insulin release and glucose metabolism, a series of experiments were conducted in TET1 and TET3 deficient mice, and it was found that there were no significant differences in glucose tolerance and insulin release between TET1/TET3 WT and KO mice, suggesting that TET2 can specifically affect the function of islets and lead to the disturbance of glucose metabolism in mice with age.

In addition, to explore the potential mechanisms of improved glucose metabolism and increased insulin secretion in TET2 deficient mice, we isolated islets from TET2 WT and KO mice on a Normal chow diet (NCD) for RNA sequencing and found that the expression of key beta cell identity genes was enhanced in TET2 KO mice. The expression of genes related to aging was decreased. In addition, under HFD conditions, the expression of β cell markers in the islets of TET2 KO mice was increased, while the expression of aging markers was significantly decreased, indicating that TET2 loss can inhibit β cell aging. To further analyze the effect of TET2 on β cell function, INS-1 cells overexpressed with TET2 OE were constructed, and the results showed that TET2 OE decreased the expression level of β cell identity genes, while the expression level of aging markers increased. These results suggest that TET2 promotes β cell senescence.

PTEN plays an important role in maintaining chromatin and genome stability. In-depth study of its up-down regulation mechanism can solve scientific problems from multiple dimensions and help clarify the functional mechanism of aging of T2DM islet beta cells. The role of PTEN in beta cell senescence is different from its role as a tumor suppressor gene in tumors. In pancreatic beta cells, the loss of PTEN leads to the up-regulation of the PI3K/AKT pathway and induces the proliferation capacity of beta cells (Stiles et al., 2006). Loss of PTEN prevents a decline in the proliferative ability of older beta cells and restores the ability of older beta cells to respond to damage induced regeneration (Zeng et al., 2013). In addition, there is growing evidence that PTEN plays an important role in the nucleus through protein-protein interactions. Cells lacking PTEN are very sensitive to DNA damage (Bassi et al., 2013). The important role of the C-terminal of PTEN in maintaining genomic stability (Sun et al., 2014). The interaction between PTEN and histone acetyltransferase is particularly evident in the control of chromatin dynamics and global gene expression (Yang & Yin, 2020)]. In addition, the interaction of PTEN with histone H1 suggests that this powerful gene may influence the expression of thousands of genes and the degree of chromatin concentration (Chen et al., 2014). Our previous studies found that the PTEN promoter region showed a low methylation level in T2DM population, suggesting that PTEN promoter methylation is closely related to the occurrence and development of T2DM (Yin et al., 2018). However, the overall effect of TETs on DNA methylation in beta cells and whether it regulates PTEN promoter methylation has not been clarified. To assess the overall effect of TETs deletion on DNA methylation, genome-wide bisulfite sequencing was performed in HG culture INS-1 using post-bisulfite aptamer labeling, and it was found that hypermethylated CpGs tended to cluster at promoter and enhancers sites in HG culture beta cells with TETs deletion. Gene-specific examination of methylated group data showed that PTEN promoter DNA methylation increased significantly and PTEN promoter DNA expression decreased significantly in TET2-KO INS-1 compared with control group. Bisulfite Sanger sequencing confirmed that the PTEN promoter region showed a low methylation level in WT HG cultured INS-1 cells, which was consistent with the results of previous studies, while the DNA methylation level of KO INS-1 cells was significantly increased. Moreover, the production of 5hmC in these regions can be saved by ectopic expression of TET2 in KO INS-1. Together, these data demonstrate that TET2 loss leads to hypermethylation of the PTEN promoter region, which may reduce its expression and associated phenotypes.

Histone acetylation is one of the most common modifications of histone tails, which regulates the accessibility of DNA to various transcription factors to control gene expression (Graff & Tsai, 2013). Histone H4 Lys-16 acetylation (H4K16Ac) is the most common acetylated lysine modification, which regulates chromatin structure by acting as a switch from an inhibited state to a transcriptional active state (Shogren-Knaak et al., 2006). Histone acetyltransferase and deacetylase control H4K16ac balance. MOF (KAT8 or MYST1) is the main acetyltransferase responsible for H4K16ac (Conrad et al., 2012). Studies on mammalian cells have shown that the loss of MOF and the consequent overall reduction of H4K16ac can lead to inadequate DNA damage response, reduced fatty acid oxidation and mitochondrial respiration, cell death and apoptosis, etc. (Khoa et al., 2020). MOF regulates the differentiation and function of α cells by acetylating H4K16ac and H4K16ac binding to Pax6 and Foxa2 promoters, and its inhibition may be a potential intervention target for the treatment of T2DM (Guo et al., 2021). Studies have shown that MOF is implicated in the regulation of glucose and amino acid balance throughout the body, and it is believed that reduced MOF levels predispose animals to metabolic syndrome with common features of T2D(Pessoa Rodrigues et al., 2021). The correlation of H4K16Ac is not limited to the regulation of DNA damage repair (McGovern & Cope, 1991). Studies have found that changes in the overall deacetylation level of H4K16 are also related to aging, which can promote DNA compression in aging cells and thus exert far-reaching regulatory effects on DNA transcription (Contrepois et al., 2012). A decrease in H4K16Ac was also observed in a premature aging mouse model (Krishnan et al., 2011). In addition, H4K16ac levels in Alzheimer’s patients are significantly reduced near genes associated with linking aging and disease (Nativio et al., 2018) .

In the regulation of gene expression, there is a complex and close relationship between DNA methylation and histone modification. A comprehensive understanding of this interrelationship is lacking. Studies have shown that methylation of histone H3K9 can guide DNA methylation (Tamaru & Selker, 2001). On the other hand, DNA methylation may also affect histone methylation (Jones et al., 1998). Our previous study found that the overall level of H4K16ac in T2DM population decreased significantly, suggesting that H4K16ac may play an important role in the occurrence and development of T2DM (Yin et al., 2022). In preliminary experiments, H4K16ac ChIP-seq analysis was performed on HG-cultured INS-1, MIN6, and βTC6 cell lines, and H4K16ac was found to be abundant and widespread throughout the genome. In addition, to analyze the changes of H4K16ac modification after OE TETs, TETs OE was subjected to H4K16ac ChIP-seq in INS-1 cells. The results showed that TET2 OE caused widespread deletions of H4K16ac on a genome-wide scale, while TET1 and TET3 OE did not undergo such changes. These results suggest that TET2 attenuates the overall modification level of H4K16ac in beta cell lines.

In order to further investigate the mechanism of TET2 promoting β cell senescence, we analyzed the RNA-seq data of TET2−/− mouse islets and found that MOF was significantly upregulated in TET2−/− mice. Further detection of MOF expression in INS-1 cells showed that OE TET2 inhibited MOF expression, while H4K16ac modification was down-regulated, which was consistent with the results of pre-test H4K16ac ChIP-seq. Previous studies have shown that PTEN physically interacts with H1, inhibiting MOF binding chromatin and reducing H4K16ac levels (Sun et al., 2014). Accordingly, we analyzed the expression of PTEN and beta cell identity genes in TET2 OE INS-1 cells, and found that PTEN increased with the decrease of beta cell specific genes. In addition, shPTEN reversed the changes in MOF and Pdx1 induced by TET2 overexpression.

To further determine whether MOF pathway is involved in β cell senescence, by inducing MOF expression in INS-1 cells, it was found that p16 was significantly inhibited after MOF overexpression. MOF OE salvaged the expression of MOF in TET2 OE cells, and found that the expression level of aging markers was significantly reduced, while the level of key identity genes in beta cells was significantly increased. These results were confirmed in MOF and TET2 dual OinS-1 cells. In addition, by constructing TET2 knockdown INS-1 cells, it was confirmed that when TET2 knockdown was performed, the MOF signaling pathway was enhanced, the expression of aging markers was reduced, and insulin secretion was increased. These preliminary results suggest that TET2-mediated PTEN demethylation and epigenetic modifications such as H4K16ac may promote β cell senescence.

In conclusion, based on the above prediction, we clarified the regulatory relationship between the molecules of the TET2/PTEN/MOF axis from the cellular and animal levels, and identified the key genes of H4K16Ac modification mediated by the TET2/PTEN/MOF axis that affect the aging of pancreatic β cells. In order to confirm the new mechanism that TET2-mediated PTEN promoter methylation can drive T2DM pancreatic β cell senescence by regulating H4K16ac modification, and provide a new drug target and theoretical basis for T2DM β cell senescence treatment.

## AUTHOR CONTRIBUTIONS

Conceptualization, W.C., and L.Y.; data curation, Q.S., H.L., X.M., Y.S., W.C., and L.Y.; formal analysis, W.C., and L.Y.; funding acquisition, W.C., and L.Y.; investigation, W.C., and L.Y.; methodology, W.C., and L.Y.; project administration, W.C., and L.Y. software, W.C., and L.Y.; supervision, W.C., and L.Y.; validation, W.C., and L.Y.; writing–original draft, W.C.; writing– review and editing, W.C., and L.Y. All authors have read and agreed to the published version of the manuscript.

## DATA AVAILABILITY STATEMENT

All analytical data were available upon request to the lead contact (L. Yin.).

